# Pre-treatment of the clinical sample with Proteinase K allows detection of SARS-CoV-2 in the absence of RNA extraction

**DOI:** 10.1101/2020.05.07.083139

**Authors:** Larissa Mallmann, Karoline Schallenberger, Meriane Demolliner, Ana Karolina Antunes Eisen, Bruna Saraiva Hermann, Fágner Henrique Heldt, Alana Witt Hansen, Fernando Rosado Spilki, Juliane Deise Fleck

## Abstract

COVID-19 (Coronavirus Disease 2019) outbreak was declared a pandemic, by World Health Organization, on March 11, 2020. Viral detection using RT-qPCR has been among the most important factors helping to control local spread of SARS-CoV-2 and it is considered the “gold standard” for diagnosis. Nevertheless, the RNA extraction step is both laborious and expensive, thus hampering the diagnosis in many places where there are not laboratory staff of funds enough to contribute for diagnosis efforts. Thus, the need to simplify procedures, reduce costs of the techniques used, and expand the capacity of the number of diagnostics of COVID-19 is imperative. In this study, detection of SARS-CoV-2 in the absence of RNA extraction has been successfully achieved through pre-treatment of the clinical sample with Proteinase K. The results show that only the use of proteinase K, without the need to perform the whole standard protocol for sample extraction and purification, can be an efficient technique for the diagnosis of COVID-19, since 91% of the samples matched the results with the standard procedure, with an average increase of 5.64 CT in the RT-qPCR.

## Introduction

The SARS-CoV-2 (Severe Acute Respiratory Syndrome Coronavirus 2) is a new human coronavirus responsible for the 2020 pandemic, causing the 2019 Coronavirus Disease (COVID-19). The virus was discovered in December 2019, in Wuhan, Hubei province, Chine, where adults were suffering from pneumonia without a known pathological agent (SINGHAL, 2020). The clinical symptoms of this infection are nonspecific, but the most common are fever, fatigue, coughing, and myalgia, like Flu symptoms, which can evolve into a SARS condition, or can be an asymptomatic disease (ZU et al., 2020). The virus rapidly spread from Wuhan to the World, infecting 3.595.662 individuals and causing the death of 247.652 in more than 215 territories until May 5th (WHO, 2020).

The relevance and importance of mass diagnosis to find the asymptomatic individuals is widely recognized as a mandatory tool to reinforce the control measures to monitor the circulation of the virus to avoid the spread of SARS-CoV-2. Molecular biology for viral detection using RT-qPCR is considered the “gold standard”. However, it is a high-cost and labor technique, that requires time and skilled personnel.

The RNA extraction step is sometimes both more laborious and expensive, thus hampering the diagnosis in many places where there are not laboratory staff of funds enough to contribute for diagnosis efforts. The use of several reagents and commercial kits and the high demand for SARS CoV-2 diagnostics, which occurs worldwide, can induce a lack of necessary chemical materials. The *American Society for Microbiology* has already expressed concern about the shortage of reagents for COVID-19 testing (AKST, 2020) as well as many other scientific societies and governments. Therefore, the need to simplify procedures, reduce the cost of the techniques used, and expand the capacity of the number of diagnostics of COVID-19 is extremely important, especially in countries and geographic regions passing through shortage of funding and supplies. In order to optimize the diagnosis process, three strategies of direct RT-qPCR (in the absence of RNA extraction) were evaluated in clinical sample positive SARS-CoV-2, previously detected with the standard procedure, i.e. extracted with commercial MagMAX™ CORE Nucleic Acid Purification Kit (Thermo Fisher Scientific).

## Material and Methods

Clinical specimens were collected with nasopharyngeal and oropharyngeal swabs submitted to routine diagnosis by the local municipalities from suspected patients (n=23) living in the Vale dos Sinos region, Southern Brazil. All samples were delivered under refrigeration to the Molecular Microbiology Laboratory (Feevale University, Novo Hamburgo, Brazil) eluted in 2 mL of saline solution, following CDC (USA) guidelines for sampling of respiratory specimens. Personal data from the patients was not accessed in the present study, following standard ethical regulatory guidelines.

All samples (n=23) showed positive results for the presence of SARS-CoV-2 RNA using the standard method. In this, viral RNA extraction was performed using the commercial MagMAX™ CORE Nucleic Acid Purification Kit (Thermo Fisher Scientific). The manual extraction technique has been validated by the Molecular Microbiology Laboratory at Feevale University.

Pilot assays were conducted employing different strategies for direct RT-qPCR (in the absence of RNA extraction). These were evaluated in comparison with RT-qPCR results obtained with prior viral RNA extraction (standard procedure), as described below.

The first pilot assay was made according to Fomsgaard and Rosenstierne (2020) study. For this, four samples (3 positives and 1 negative for the standard procedure) were tested. The samples were analyzed pure and diluted, 1:1 with PBS, and heated to 98° and 70°C for 3 and 5 minutes, followed by incubation at 4°C until the time of analysis, totaling eight trials to each sample (FOMSGAARD; ROSENSTIERNE, 2020).

In the second pilot assay, one positive sample was submitted to twelve trials. The tests were performed to 5 and 8 min, at 70 ° and 98 ° C, followed by incubation at 4°C until the time of analysis. The sample was tested pure, combined with Proteinase K, and with Lysis Buffer.

Analyzing that the best strategy was to combine the sample with proteinase K (PK), a third test was performed. Twenty-three positive samples were combined with PK, respecting a 1:21 ratio of PK to each sample. After, the sample with PK was submitted to 98°C for 5 min, followed by incubation at 4°C until the time of analysis.

For the detection of SARS-CoV-2, the Reverse Transcriptase technique was used followed by the Polymerase Chain Reaction (RT-qPCR) in one step, using the E gene target from the Charité protocol (CORMAN et al., 2020). The reaction was done with 45 cycles and using 10μL of Buffer and 0.8 μL of 25x Enzyme by AgPath-ID™ One-Step RT-PCR Kit, and 0.8 μL of each primer (R and F) and 0.4 μL of probe P1, from Applied Biosystems. The DNAse/RNAse Free Water was 2.2 μL for the procedure standard reactions, which use 5μL of sample, and 0.2 μL for the three strategies, that use 7 μL of sample.

## Results and Discussion

The first pilot assay showed that heat the pure samples for 5 minutes at the both temperatures had been the best parameter. The results showed 87.5% and 62.5% of agreement between the standard treatment using heat-treatments of 70° and 98°C, respectively. The results also showed that diluting the sample with PBS is not ideal, on average, the treatments showed only 33.33% of the relation to the standard treatment. Thus, a second strategy was proposed.

In the second strategy, heating of the samples combined with PK to 98°C, SARS-CoV-2 was detected at both times (5 and 8 minutes), and cycle threshold (CT) values were next to standard procedure. Otherwise, there was not detection with heating to 70°C, just like in the trials where Lysis Buffer was added. Considering that the best result of the second strategy was to use PK, the third test was performed using it and incubating for five minutes at 98°C.

For the third test, the strategy showed that of the twenty-three samples analyzed, only two were not detected. This represents 91% of agreement with standard procedure, presenting an average increase of 5.64 CT in the RT-qPCR. The non-detection of the two samples can indicate the presence of RT-qPCR inhibitors. Both samples showed at 37.8 CT, in standard procedure. The CT mean of the results is expressed in table 1, with a mean standard deviation of 2.77.

**Table 1:**
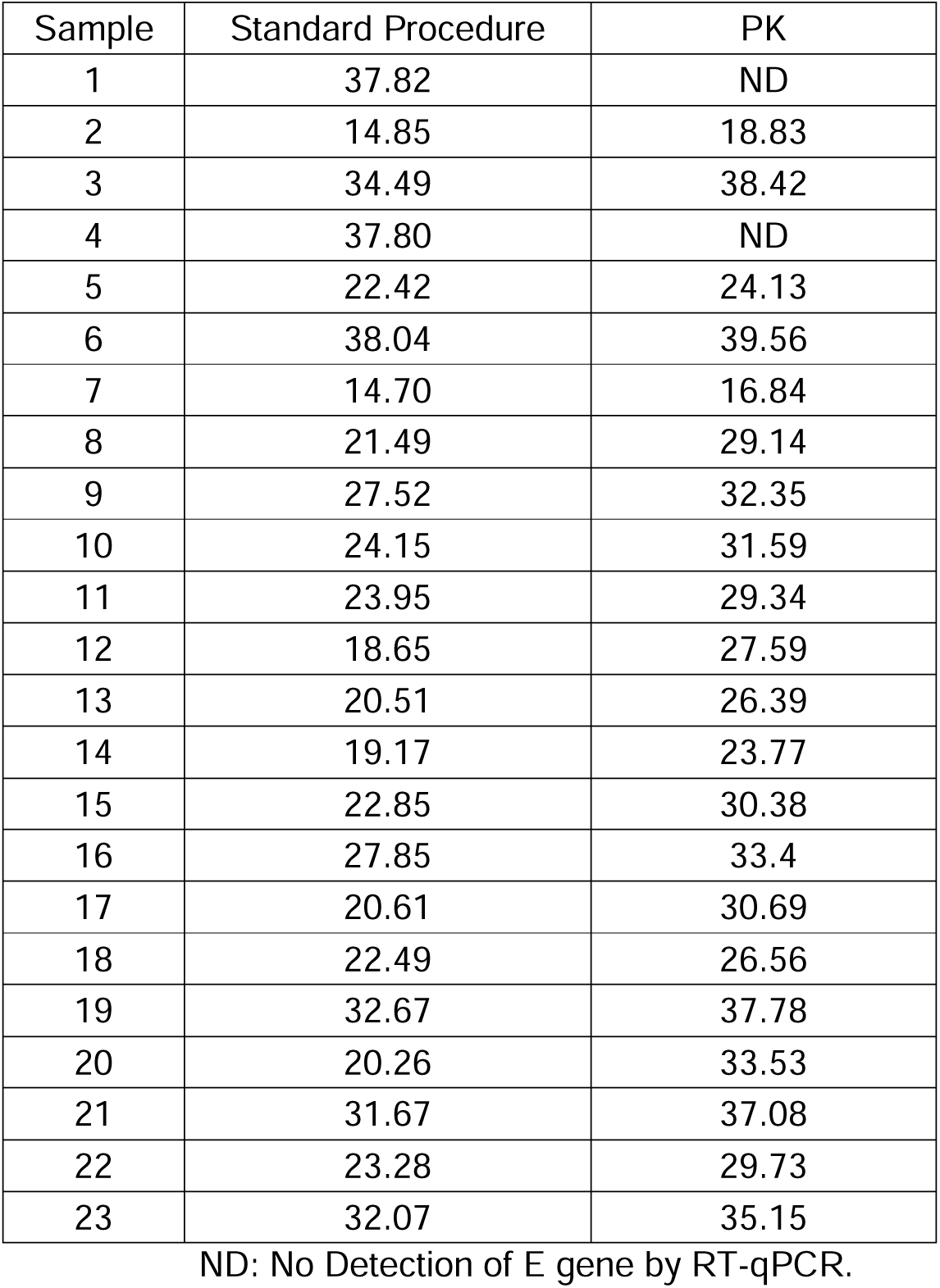
CT mean from the PK pretreatment in comparison as standard procedure.

| Sample | Standard Procedure | PK |
| --- | --- | --- |
| 1 | 37.82 | ND |
| 2 | 14.85 | 18.83 |
| 3 | 34.49 | 38.42 |
| 4 | 37.80 | ND |
| 5 | 22.42 | 24.13 |
| 6 | 38.04 | 39.56 |
| 7 | 14.70 | 16.84 |
| 8 | 21.49 | 29.14 |
| 9 | 27.52 | 32.35 |
| 10 | 24.15 | 31.59 |
| 11 | 23.95 | 29.34 |
| 12 | 18.65 | 27.59 |
| 13 | 20.51 | 26.39 |
| 14 | 19.17 | 23.77 |
| 15 | 22.85 | 30.38 |
| 16 | 27.85 | 33.4 |
| 17 | 20.61 | 30.69 |
| 18 | 22.49 | 26.56 |
| 19 | 32.67 | 37.78 |
| 20 | 20.26 | 33.53 |
| 21 | 31.67 | 37.08 |
| 22 | 23.28 | 29.73 |
| 23 | 32.07 | 35.15 |

ND: No Detection of E gene by RT-qPCR.

Comparing our first pilot assay with Fomsgaard and Rosenstierne’s study, the results showed that incubation at 98°C for 5 minutes is the best trial for the two studies, although that the number of samples tested was bigger (37 positives and 22 negatives).

The study of Merindol et al. (2020), which compares forms to stored samples, showed that it is recommended the RNA extraction for stored samples with saline solution, where a loss of 10 CTs were observed. Otherwise, in this study, all samples are stored with saline solution and occurred the detection of 91% of them, by PK pretreatment and direct RT-qPCR (in the absence of RNA extraction). Our findings showed consistently that proteinase K pre-treated samples may be readily usable for RT-qPCR under emergency situations and population studies, in situations where there is momentary lack or shortage of reagents for viral RNA extraction.

